# Protein accumulation upstream of Cdk1 activation regulates the competence and the timing of meiosis resumption

**DOI:** 10.64898/2026.06.25.734491

**Authors:** Mariam Gachechiladze, Sarah B. Eivers, Robert Poulhe, Matthew Sonnett, Leonid Peshkin, Catherine Jessus, Enrico Maria Daldello

## Abstract

The capacity to resume meiosis is progressively acquired during oogenesis and is ultimately restricted to fully grown oocytes. Meiotic resumption is triggered by hormonal stimulation and requires activation of Cdk1, the universal driver of M-phase entry. Cdk1 activation occurs in two steps: an initial activation of Cdk1, followed by an amplification phase that drives cell cycle re-entry. The first step depends on the accumulation of proteins that promote Cdk1 activation, while the second step involves a regulatory network of kinases and phosphatases. Using TMT-based quantitative proteomics, we reveal that growing oocytes first acquire the ability to regulate protein homeostasis in response to hormonal stimulation, and only later gain the competence to amplify initial Cdk1 activity and enter meiosis. Notably, protein accumulation, occurring independently of Cdk1 activation, is controlled by both translational and non-translational mechanisms. Together, our findings show that the molecular competence to trigger Cdk1 activation is acquired in a stepwise manner during oocyte growth. The earliest regulatory layer is the acquisition of the ability to respond to hormonal stimulation by accumulating proteins that are required for efficient Cdk1 activation and meiotic resumption.

## INTRODUCTION

In all metazoans, oocytes arrest for a long period of time in prophase of the first meiotic division, allowing the oocyte to undergo a sustained growth phase in which it accumulates maternal RNAs, proteins, and organelles necessary to support both meiotic progression and early embryogenesis^1,2^. The oocyte growth phase is a critical determinant of developmental competence. At the end of the growth phase, oocytes must acquire the ability to resume meiosis and to execute the two meiotic divisions at the time of ovulation. A tight temporal regulation of the competence acquisition is important to prevent small oocytes to be able to resume meiotic divisions^3^. In *Xenopus laevis* full-grown oocytes, a hormonal stimulation, progesterone, triggers the release of the prophase block, whereas small growing oocytes are unable to resume meiosis in response to the hormone. In fully-grown oocytes, progesterone produces a decrease in cAMP and PKA activity, which promotes a change in protein homeostasis resulting in the translation and accumulation of proteins that are required for the initial activation of Cdk1^4^. Cdk1 activation then engages a series of feedback loops leading to its full activation, known as the autoamplification loop. Indeed, Cdk1 directly and indirectly activates the Cdc25 phosphatase, which in turn removes inhibitory phosphates from Cdk1, while simultaneously inhibiting Myt1, the kinase responsible for Cdk1 inactivation^5^. This mutually reinforcing network creates a bistable switch ensuring robust and irreversible entry into meiotic M-phase. Additional components are involved in the feedback loop, such as the Polo-like kinase Plk1, which contributes to Cdc25 activation, Bora, a cofactor that promotes Plk1 activity^6^, Aurora-A ^7,8^ and the Gwl-Arrp19 axis, which inhibits PP2A phosphatase that opposes Cdk1 substrate phosphorylation^9^. Consistent with the importance of translation in initiating Cdk1 activation, blocking protein synthesis with cycloheximide (CHX) prevents Cdk1 activation in all vertebrates studied to date, including *Xenopus* and humans, with the notable exception of small rodents^10,11^. These findings demonstrate that proteins must accumulate in response to progesterone to trigger the activation of Cdk1. Some of these proteins have been identified, including Mos^12,13,11^, Cyclin B1^14,15,11^, and Speedy/RINGO^16,17^. Mos is an oocyte specific kinase whose translation is activated by progesterone^12,13,11^. It triggers the MAPK cascade, which contributes to the activation of Cdc25, promoting the conversion of stockpiled Cdk1/Cyclin B2 complexes that can activate the autoamplification loop. In addition, Mos is also required to prevent DNA replication during the MI-MII transition and to maintain Cdk1 activity during metaphase II arrest^18–21^. Cyclin B1 is a direct activator of Cdk1, which accumulates following progesterone treatment and binds to free Cdk1, forming active Cdk1/Cyclin B1 complexes that can initiate the autoamplification loop. Speedy/RINGO is also an activator of Cdk1 that can bind directly and activate the kinase independently of Cyclin B. Speedy/Cdk1 complexes could also provide the initial activity required to launch the autoamplification loop^22^. The presence of multiple partially redundant pathways likely reflects the evolutionary pressure to ensure robust activation of Cdk1 and the fidelity of meiotic re-entry^23^. This redundancy may also indicate that additional, yet unidentified components contribute to linking PKA downregulation to Cdk1 activation, highlighting the importance of exploring the upstream regulatory network and identify its components. In support of this idea, preventing the accumulation of known upstream components results in a delay in Cdk1 activation, suggesting that these pathways act in parallel and partially compensate for one another to guarantee timely meiotic resumption^4,19^.

A major challenge lies in understanding how oocyte growth is coordinated with the multistep pathway required for Cdk1 activation. These mechanisms are essential to connect meiotic resumption with ovulation, an event that must occur exclusively in fully-grown oocytes. Yet how and when the molecular ability to initiate these steps is acquired during oogenesis remains unclear. A second challenge is to identify additional proteins that accumulate before the first activation of Cdk1 and that functionally link PKA downregulation to Cdk1 activation. However, translation is activated both before and after Cdk1 activation. For this reason, most large-scale studies have not distinguished between upstream (Cdk1-independent) and downstream (Cdk1-dependent) changes in protein abundance^24–27^. This distinction is essential to elucidate causality in meiotic resumption.

In this study, we addressed these challenges combining TMT-based mass spectrometry to our Cip1-based approach^11,15^ to define the landscape of protein abundance changes during oocyte maturation before and after Cdk1 activation. Our results reveal that the regulation of protein homeostasis, via both translation dependent and independent-mechanisms, plays a central role in shaping the molecular competence for Cdk1 activation and highlights the developmental windows during which the oocyte progressively acquires the components of the molecular pathway required to resume meiosis.

## RESULTS

### The competence to activate Cdk1 is acquired progressively during oocyte growth

*Xenopus* growing oocytes have been classified in 6 categories (I to VI) based on diameter and pigmentation criteria^1^. Full-grown oocytes (diameter≥1200µm) are responsive to progesterone, the hormone that triggers meiotic maturation in *Xenopus* oocytes. Progesterone triggers a drop of adenylate cyclase activity, and consequently of cAMP and PKA activity that is both necessary and sufficient to allow Cdk1 activation, through the translation of new proteins, as Mos, and the stabilization of Cyclin B1. In contrast, stage IV oocytes (ranging from 600 to 1000µm) are unable to activate Cdk1 and to resume meiosis in response to progesterone^28,29^. Nevertheless, these small oocytes respond to hormone with a decrease in adenylate cyclase activity and in their cAMP levels, suggesting that stage IV oocytes are unable to activate events required for Cdk1 activation that occur downstream PKA inactivation^30^. Therefore, we wondered whether Mos translation and Cyclin B1 accumulation take place in stage IV oocytes stimulated by progesterone^11,15,31^. As stage IV oocytes span a wide diameter range with varying pigmentation and morphology, we collected oocytes grouped by 100µm intervals (Fig. 1A). Each group included fully-grown oocytes as positive controls (Fig. 1A). Oocytes were stimulated by progesterone and 18 hours later, they were scored for the appearance of a white spot, the marker of the nuclear envelope breakdown (NEBD), *i.e.* Cdk1 activation. 100% fully-grown oocytes showed a clearly defined white spot, reflecting NEBD and Cdk1 activation, while oocytes with a diameter≤1000µm showed no pigment re-arrangement, suggesting the absence of Cdk1 activation. Oocytes around 1100µm constituted a mixed population, 30 to 40% of them showing a white spot in response to progesterone (Fig. 1A and Supp. Fig. 1A). These results are in good accordance with previous findings showing that stage IV oocytes are unable to activate Cdk1^28^. Oocytes were lysed in different volumes of buffer depending on their diameter, in order to achieve extract with equal protein concentration (Supp. Fig. 1B). *In vitro* kinase assays revealed that Cdk1 was activated only in oocytes that displayed a white spot, *i.e.* NEBD (Fig. 1B).

**Figure 1:**
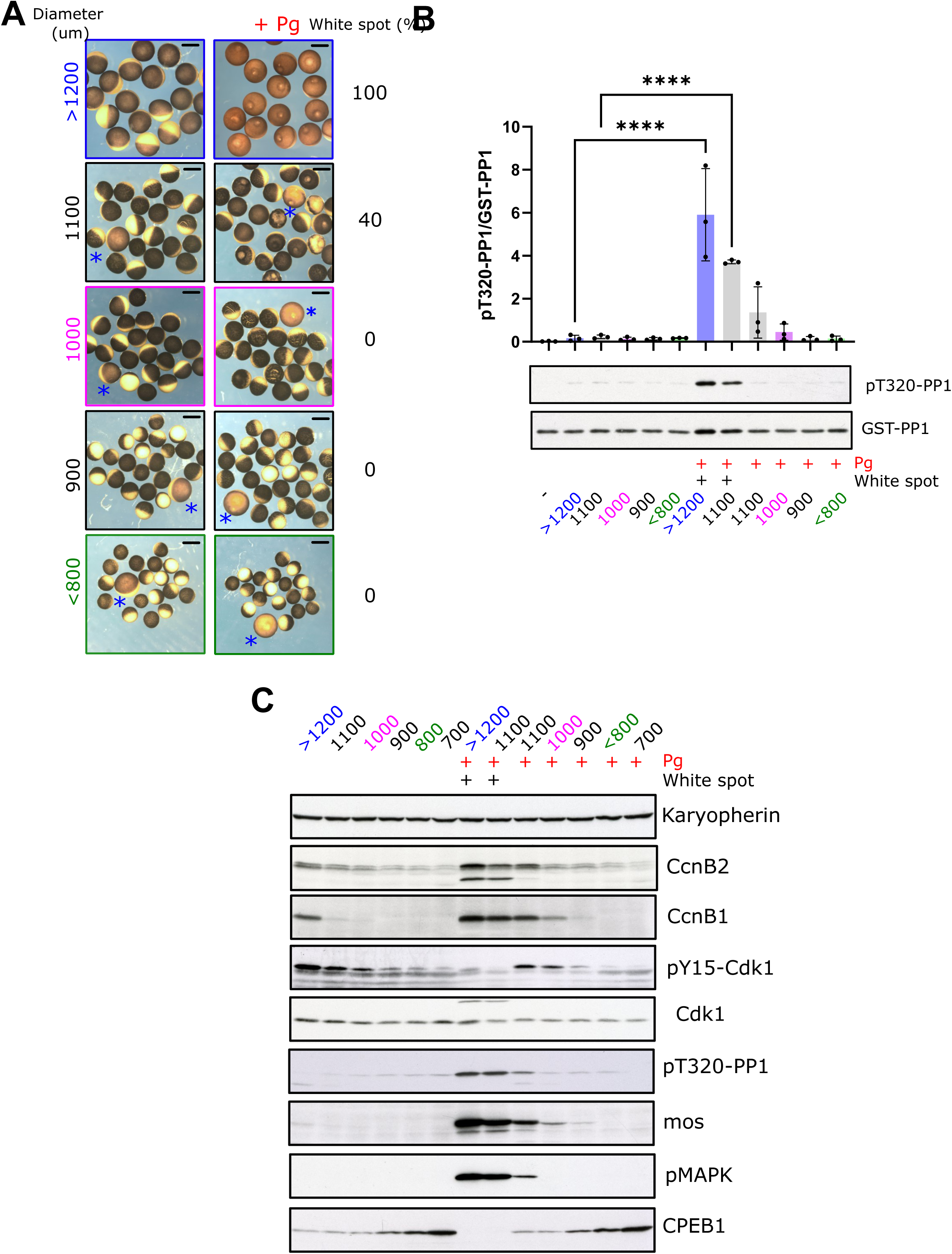
Acquisition of the ability to modulate protein homeostasis precedes full meiotic competence. (A) Oocytes were divided in different groups according to their size. Oocytes were incubated with progesterone and after 18 h picture were taken and the percentage of oocytes undergoing nuclear envelope breakdown (NEBD) was calculated by looking at the appearance of the white spot. Fully grown oocytes (>1200 µm) were used as positive controls. n = 30 oocytes per group. (B) In vitro Cdk1 kinase assay on oocyte lysates from each diameter group. Cdk1 activity was assessed by phosphorylation of recombinant GST-tagged PP1 on T320. 3 biological replicates. Statistical significance was assessed using ANOVA; ****p < 0.0001. (C) Western blot analysis of markers of meiotic maturation and protein accumulation in oocytes of increasing diameter before and after 18 hrs after progesterone stimulation.

We then determined from which diameter oocytes become capable to accumulate Cyclin B1 and translate Mos in response to progesterone, two events that occur upstream of Cdk1 activation in fully-grown oocytes^11,32,33^. As expected, Cdk1 was fully activated in the ≥1200µm oocytes, as demonstrated by the phosphorylation of PP1 at T320, which tracks the first activation of Cdk1, and the dephosphorylation of Cdk1 at Y15, which marks the activation of the autoamplication loop (Fig. 1C). Mos and Cyclin B1 accumulated in these oocytes and MAPK was phosphorylated as a consequence of Mos expression (Fig. 1C). Surprisingly, in response to progesterone, Cyclin B1 and Mos accumulate in the 1000µm oocytes (Fig. 1C), although these oocytes neither display a white spot nor support Cdk1 activation, as shown by the maintenance of Cdk1 phosphorylation at Y15 (Fig. 1B-C). As expected, the 1100µm group splits into two populations. The 1100µm-oocytes with a white spot activate Cdk1, as shown by Cdk1 dephosphorylation at Y15 and the phosphorylation of PP1 (Fig. 1C). This group of oocytes also accumulate Cyclin B1 and Mos and they activate MAPK (Fig. 1C), similarly to what is observed in fully-grown oocytes. In contrast, 1100µm oocytes that do not display a white spot are comparable to 1000µm oocytes: they accumulate Cyclin B1 and Mos, and even support weak MAPK activation, but they are unable to fully activate Cdk1, as indicated by the maintenance of its phosphorylation on Y15 (Fig. 1C). Hence, 1000µm and 1100µm oocytes that are unable to undergo NEBD in response to progesterone are characterized by their ability to accumulate Mos and Cyclin B1, although this is not sufficient to trigger Cdk1 activation. Finally, <800um oocytes are not able to support the activation of Cdk1 nor to accumulate Cyclin B1 or Mos (Fig. 1C). Interestingly, the level of Cyclin B2 is very low in <800um oocytes, progressively increasing during the oocyte growth (Fig.1C). Accordingly, Y15-phosphorylation of Cdk1, which takes place when Cyclin B binds to Cdk1, increases proportionally with the amount of Cyclin B2 during oocyte growth (Fig. 1C). This result indicates that ≤800um oocytes have less Cdk1-Cyclin B2 complexes, which accumulate progressively during the growth to form the well-characterized pre-MPF molecules characterizing the end of the G2-phase, formed of Cdk1-phosphorylated at Y15 bound to Cyclin B2.

Three distinct diameter groups can thus be distinguished based on their pre-MPF content and their response to progesterone: the ≥1200µm oocytes are equipped with Cdk1 phosphorylated at Y15-Cyclin B2 (pre-MPF), progesterone triggers accumulation of Cyclin B1 and Mos as well as stimulation of protein translation, which leads to full activation of Cdk1; the 1000µm oocytes also contain pre-MPF and progesterone stimulates Cyclin B1 and Mos accumulation but does not lead to Cdk1 full activation; finally, the ≤800µm oocytes with low level of pre-MPF, progesterone is incapable of inducing Cyclin B1 or Mos accumulation and therefore cannot activate Cdk1.

### Changes in protein abundance induced by progesterone upstream Cdk1 activation in intermediate and full-grown oocytes

In fully-grown oocytes, Cyclin B1 accumulation and Mos translation are induced by progesterone upstream of Cdk1 activation and are required for this activation^11,13,14,23^. In contrast, in 1000µm oocytes, progesterone does not promote Cdk1 activation, although it triggers elements of the upstream signaling cascade, such as Cyclin B1 accumulation and translational activation (Fig. 1C-D). Therefore, we aimed to assess whether the global repertoire of proteins accumulated in response to progesterone prior to Cdk1 activation was comparable between 1000µm and 1200µm oocytes by conducting a TMT-mass spectrometry experiment (Fig. 2A). 800µm oocytes were used as negative control as they do not support Cyclin B1 accumulation and translational activation in response to progesterone (Fig. 1C). In 1200µm oocytes, progesterone induces two waves of protein accumulation: the first, which is the focus of this study, occurs prior to Cdk1 activation, and the second occurs thereafter^11^. To compare the proteins involved in the first wave between 1000µm and 1200µm oocytes, it was necessary to exclude those associated with the second wave. For this purpose, we used the Cip1 protein, a direct inhibitor of Cdk1 activity^11,15^. The Cip1-based approach blocks Cdk1 activation in fully-grown oocytes, enabling the analysis of progesterone-induced events independently of Cdk1 activity^11,15^ (Fig. 2A). We confirmed that Cip1 injection prevents Cdk1 activation, as ascertained by Cdk1 dephosphorylation at Y15 and by *in vitro* kinase assay (Supp. Fig. 2). Importantly, molecular events triggered by PKA downregulation that occur upstream of Cdk1 activation, such as Cyclin B1 accumulation, are still induced by progesterone in Cip1-injected oocytes^11,15^ (Supp. Fig. 2A). In contrast, events known to occur downstream of Cdk1 activation, such as MAPK activation, CPEB1 degradation and S287 dephosphorylation of Cdc25, do not take place in Cip1-injected oocytes^11^ (Supp. Fig. 2A). Our TMT-mass spectrometry experiment quantified the abundance of over 240,000 peptides corresponding to 12,000 proteins (Dataset 1). Taking peptide coherence in account (see appendix A), 213 proteins accumulated, while 177 decreased during the transition from prophase (PRO) to metaphase I (MI) triggered by progesterone in fully-grown oocytes (LogFC +/- 0.4, FDR<0.05, and peptide coherence >60%) (Supp. Fig. 3A). Interestingly, in fully-grown oocytes injected with Cip1, thus without Cdk1 activation, 65 proteins increased and 21 decreased in response to progesterone (Supp. Fig. 3B). As expected, minimal changes were induced by the injection of Cip1 in prophase-arrested oocytes (Supp. Fig. 3C). In 1000µm oocytes, the extent of progesterone-induced changes in protein abundance, 132 increasing and 127 decreasing, was intermediate between that observed in fully-grown oocytes with or without Cdk1 activation (Supp. Fig. 3D). As anticipated, only minor changes were detected in 800µm oocytes (Supp. Fig. 3E).

**Figure 2:**
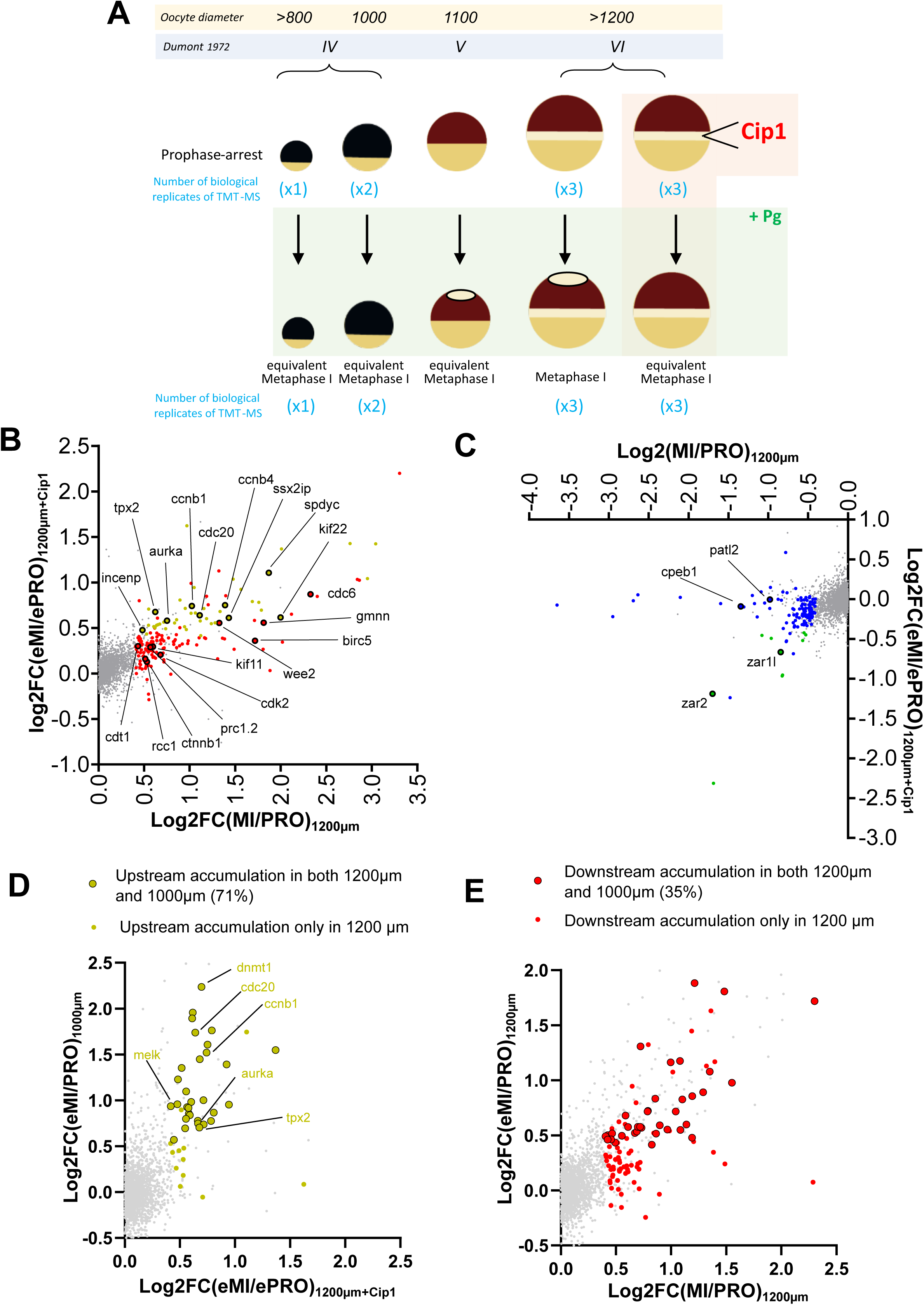
Two waves of protein accumulation take place during meiosis resumption. (A) scheme of the oocytes and the conditions (18 samples) used in the TMT-mass spectrometry experiment. (B-C) Scatter plot comparing log₂ fold changes between MI/PRO (x-axis) and eMI/ePRO (y-axis). The names of proteins that were previously reported to accumulate (B) or be degraded (C) during meiotic maturation from prophase I to metaphase II are highlighted. The supporting references are provided in Supplementary Figure 4. (D–E) Scatter plots comparing protein abundance changes in 1000 µm oocytes after progesterone treatment (x-axis) with changes observed in fully grown oocytes before Cdk1 activation (eMI/ePRO, y-axis). (D) Early-upregulated proteins (upstream of Cdk1). (E) Late-upregulated proteins (Cdk1-dependent). Color-coding distinguishes proteins whose accumulation is detected or absent in 1000 µm oocytes.

**Figure 3.**
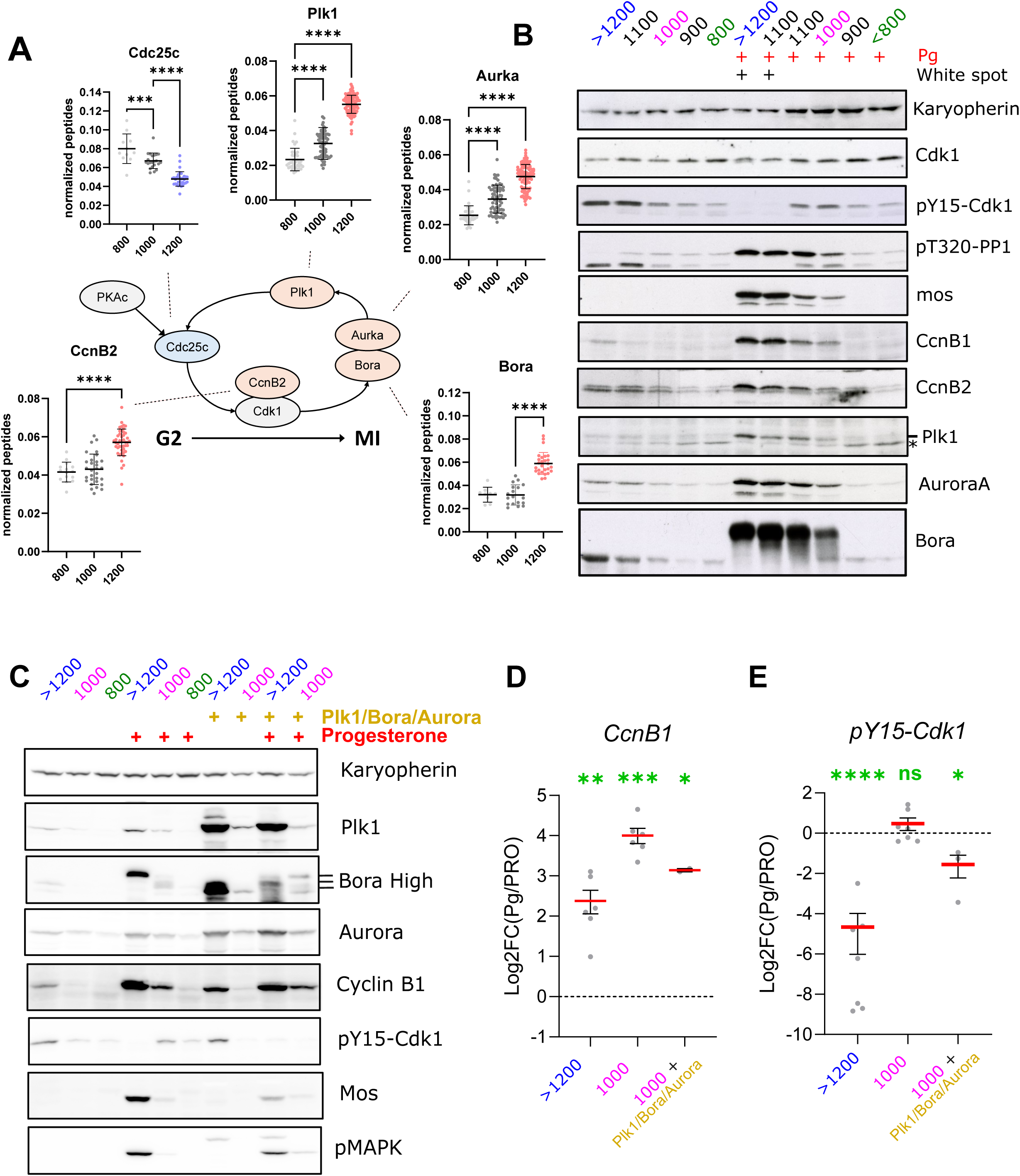
Key regulators of meiotic resumption accumulate during the final stages of oocyte growth. (A) Normalized peptide abundance of selected components of the Cdk1 activation network in oocytes of increasing diameter (800 µm, 1000 µm, 1200 µm), as measured by TMT-based quantitative proteomics. Each dot represents a peptide; bars indicate the mean. Statistical significance was assessed by ANOVA with ***p < 0.001. ****p < 0.0001. (B) Western blot analysis in oocytes grouped by size and treated or not with progesterone. (C-E) Western Blot analysis in oocytes grouped by size, treated or not with progesterone and injected or not with Plk1/Aurka/Bora mRNA mix (D-E) The fold-change of the levels of Cyclin B1(D) pr pY15-CDk1 (E) between prophase and Progesterone-treated samples is plotted for 1200um; 1000um and Plk1/Aurka/Bora-injected 1000um oocytes. Statistical significance was assessed by one sample T test with *p<0.1. **p<0.01. ***p<0.0001. ****p<0.00001

To validate our dataset, we compared our results with previously reported proteins whose accumulation or degradation had been detected by western blot. During the PRO-MI transition in fully-grown oocytes, 35 out of 38 proteins with known accumulation or degradation patterns from low throughput approaches were correctly classified in our dataset (Supp. Fig. 4). Furthermore, the changes in protein abundance detected in our dataset during the PRO-MI transition in fully-grown oocytes correlated well with those reported in a recently published proteomic study of meiotic maturation in *Xenopus* oocytes^27^ (Supp. Fig. 5).

**Figure 4:**
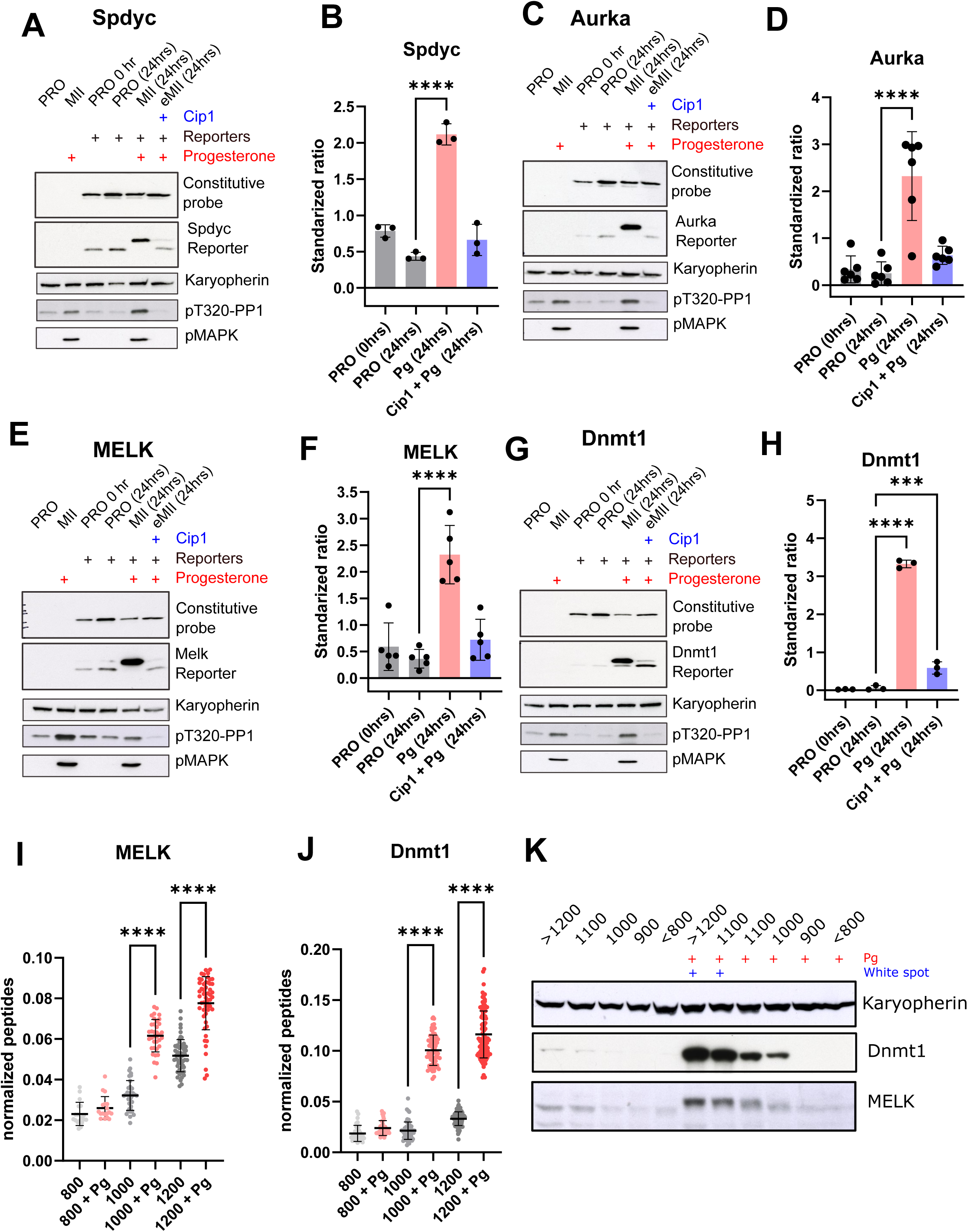
Distinct regulatory mechanisms underlie the accumulation of proteins upstream of Cdk1 activation. (A–H) Oocytes were injected with Cip1-GST and 1 hour later injected with a mix of constitutive probe and a reporter for either *Speedy/RINGO* (A-B), *Aurora A* (C-D), *Melk* (E-F), and *Dnmt1* (G-H). After an overnight incubation, oocytes were collected at prophase I (PRO, 0 h). Oocytes were treated with progesterone and collected after 24 h (MII, 24 hrs) and (eMI, 24 hrs). A control group was incubated in the absence of progesterone for 24 hrs (PRO, 24 hrs). (B-D-F-H) Quantification of standardized ratio between reporters and constitutive probes. 3 speedy (B), 6 Aurora A (D), 5 Melk (F), 3 Dnmt1 (H) biological replicates were analysed. Quantification panels show mean ± SD; statistical significance assessed by ANOVA with ***p < 0.001 and ****p < 0.0001. (I–J) Peptide-level quantification of *MELK* (I) and *Dnmt1* (J) in 800 µm, 1000 µm, and 1200 µm oocytes, with and without progesterone treatment. Values represent normalized peptide abundance across conditions. Asterisks indicate significance by ANOVA with multiple comparisons. ****p < 0.0001. (K) Western blot analysis in oocytes grouped by size and treated or not with progesterone.

**Figure 5:**
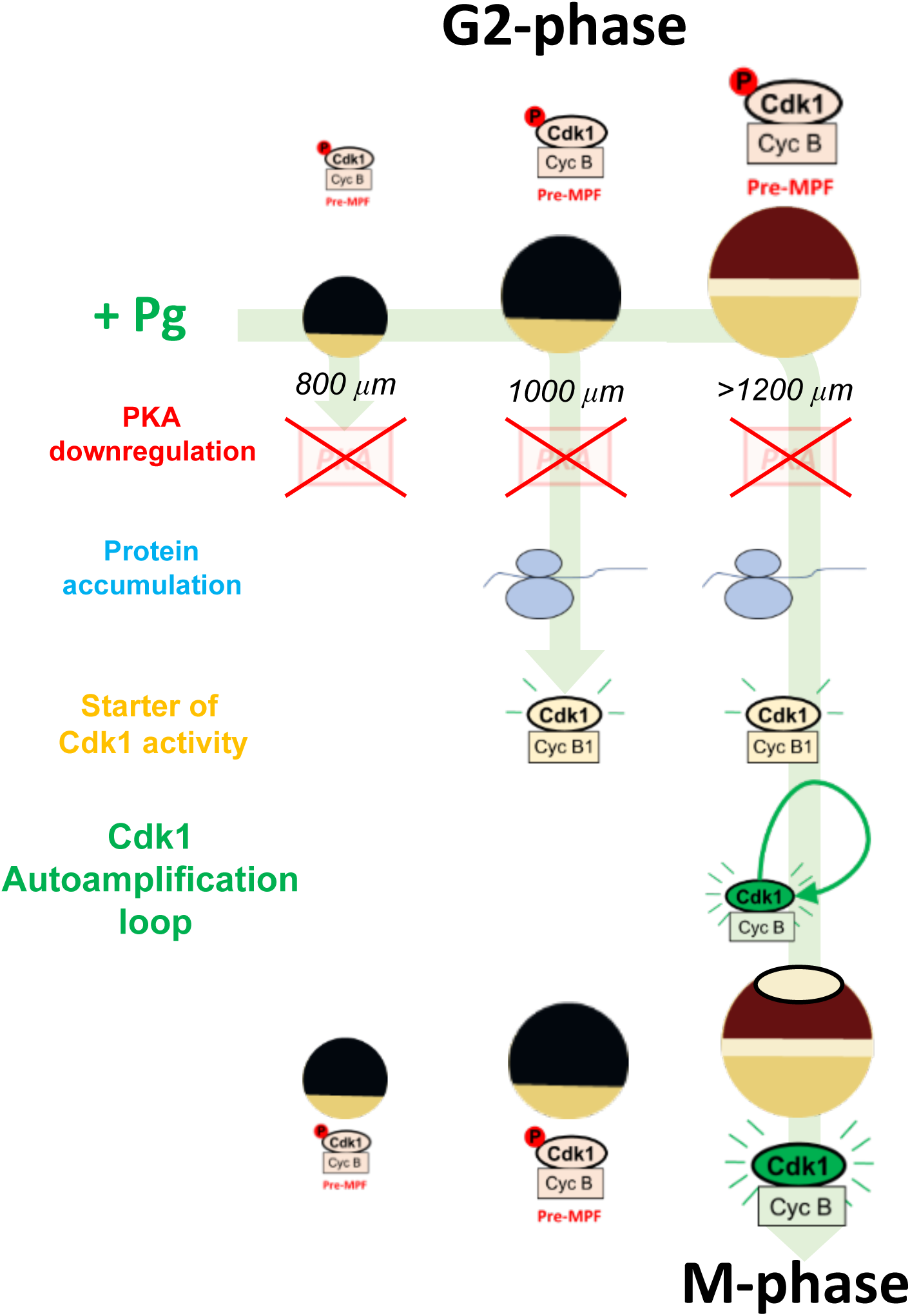
The progressive acquisition of ability to resume meiosis during Xenopus oocyte growth. Oocytes smaller than 800 µm are unable to activate protein accumulation in response to progesterone. Around 1000 µm, oocytes acquire the capacity to accumulate upstream regulators of Cdk1, but cannot yet trigger full Cdk1 activation, because of the lack of components of the autoapplication. Only fully grown oocytes (>1200 µm) are competent to enter meiosis upon progesterone stimulation, completing the two-step process of Cdk1 activation: an initial accumulation of upstream activators followed by autoamplification through feedback loops.

We next analyzed the changes in protein abundance triggered by progesterone in fully-grown oocytes, either control or Cip1-injected, to identify the changes in protein abundance occurring upstream and downstream Cdk1 activation (Fig. 2B-C). Importantly, Cyclin B1 and Aurora-A, which are known to accumulate independently of Cdk1 activation and thus represent upstream events^11,32^, and CPEB1, which is degraded downstream of Cdk1 activation and represents a downstream event^11,33^, are correctly classified in our dataset (Fig. 2B-C). Gene ontology enrichment analysis of the molecular processes and the biochemical functions among proteins, whose abundance changed during meiotic resumption, revealed a strong overrepresentation of mitotic cell cycle components (GO 1903047), kinases/phosphatases and mRNA-binding proteins (Supp. Fig. 6).

We then compared the proteins accumulating in response to progesterone in ≥1200µm oocytes and growing oocytes. Cyclin B1 and Aurora A accumulated in 1000µm oocytes but not in 800µm ones (Supp. 7A-B). We then assessed whether the fingerprint of proteins that accumulate in 1000µm oocytes in response to progesterone matches with the set of proteins that increase in abundance in fully-grown oocytes, either upstream or downstream of Cdk1 activation. While the abundance of most upstream proteins increased similarly in 1000µm and ≥1200µm oocytes, only a minority of downstream proteins accumulated in 1000µm oocytes (Fig. 2D-E). As expected, in 800µm oocytes, neither upstream nor downstream proteins accumulated in response to progesterone (Supp. Fig. 7C-D). It has been shown that the early wave of translation takes place independently of CPEB1 degradation in both *Xenopus* and mouse oocytes^11,34^. Here, we confirm that CPEB1 degradation does not occur in Cip1-injected oocytes (Fig. 2C), and showed that upstream protein accumulation still takes place under this condition, indicating that this process is not dependent on CPEB1 degradation (Fig. 2B). Interestingly, CPEB1 is more abundant in small oocytes than in fully-grown oocytes and is not degraded in response of progesterone in 1000µm oocytes (Fig. 1C). This could explain why the accumulation of downstream proteins is largely inhibited in these small oocytes, which lack Cdk1 activity and contain high CPEB1 levels.

Our results demonstrate that the competence to resume meiosis is a highly ordered process. Below 800 µm, the low level of Cyclin B and the inability to activate protein translation are the limiting factor preventing Cdk1 activation. Between 800µm and 1000µm, oocytes acquire the ability to modulate protein homeostasis in response to progesterone in a manner comparable to fully-grown oocytes. Importantly, they accumulate Cyclin B1, which in fully-grown oocytes, serves as a trigger to launch the Cdk1 auto-amplification loop. However, they remain unable to fully activate Cdk1. Between 1000µm and 1200µm, oocytes become fully competent to resume meiosis and activate Cdk1.

### Multiple components of the autoamplification loop are missing in 1000µm oocytes

In 1000μm oocytes, progesterone induces changes in protein homeostasis similar to those observed in ≥1200µm oocytes, including the formation of Cdk1-Cyclin B1 complexes. However, Cdk1 is not activated, suggesting that some components of the positive feedback network required for Cdk1 activation are missing in these oocytes. Indeed, previous work has shown that Plk1 is absent in stage IV oocytes^28^. However, its overexpression is insufficient to restore the ability of stage IV oocytes to fully activate Cdk1 in response to progesterone^28^, suggesting that additional components are missing in these oocytes. Therefore, we mined our mass spectrometry dataset to identify proteins of the Cdk1 activation network that are less abundant in small oocytes.

Our dataset confirms that Plk1 levels increase during the last steps of oogenesis and in response to progesterone (Fig. 3). Cyclin B2 abundance and Y15 phosphorylation of Cdk1 both increase with the diameter of the oocyte (Fig. 3), whereas total Cdk1 abundance remains constant (Supp. Fig. 7E). These findings indicate that the pre-MPF stockpile gradually increases toward the end of the oocyte growth period, while its level is very low in ≤800µm oocytes. Importantly, many components of the Cdk1 autoamplification loop are less abundant in small oocytes, such as Aurora-A and Bora, that are required for Plk1 activation (Fig. 3). Surprisingly, Cdc25 abundance is slightly but significantly higher in small oocytes (Fig. 3A). However, Cdc25 is fully phosphorylated at S287 in both 1000µm and full-grown oocytes (Supp. Fig. 2), which prevents its activation. This inhibitory phosphorylation is catalyzed by PKA and maintained until NEBD in progesterone-stimulated full-grown oocytes^11,35^, indicating that elevated Cdc25 levels in small oocytes cannot facilitate prophase release. Some other components of the network are instead constant in abundance (Supp. Fig. 7E-H). To validate these observations experimentally, we analyzed the abundance of these components of the Cdk1 autoamplification loop by western blotting in oocytes of different diameters. Consistent with the mass-spectrometry data, Plk1, Aurora-A and Bora levels increased during the final stages of oocyte growth (Fig. 3B). This analysis also confirmed the intermediate phenotype of 1000µm oocytes characterized by Cyclin B1 accumulation in response to progesterone and their inability to trigger Cdk1 Y15 dephosphorylation (Fig. 3B).

We next tested whether increasing the abundance of Plk1, Bora and Aurora-A was sufficient to rescue Cdk1 activation in 1000µm oocytes. Oocytes were microinjected with the three corresponding mRNAs, incubated overnight to allow protein expression, stimulated with progesterone and collected 24 hours later. In response to progesterone, Cyclin B1 accumulation took place in both 1000µm and 1200µm oocytes (Fig. 3C-D). Interestingly, co-expression of Plk1, Bora and Aurora-A rescued progesterone-induced Cdk1 Y15 dephosphorylation in 1000µm oocytes (Fig. 3C-E). This result indicates that the absence of the Bora/Aurora A/Plk1 branch of the autoamplification loop is the limiting factor for Cdk1 activation in the developing oocyte.

### Protein accumulation in response to progesterone is regulated by translational activation or protein stabilization

Given the importance of protein accumulation for Cdk1 activation, we next investigated whether these protein level changes result from increased translation or decreased degradation in fully-grown oocytes^11^. Unlike translatomic-based techniques, our proteomic approach detects protein accumulation dependent on activation of translation or inhibition of degradation. To investigate the mechanisms regulating protein accumulation during meiosis resumption, we used translational reporters, which enable to assess the translation efficiency of several proteins identified in our dataset as accumulating in fully-grown oocytes upstream of Cdk1 activation^11^. Experiments were performed using the Cip1-based approach to distinguish regulatory events taking place upstream or downstream of Cdk1 activation. We selected four proteins that accumulate upstream of Cdk1 activation and are potentially regulating cell cycle progression, Aurora-A, MELK, and Dnmt1, and studied their translational control. Aurora-A is a protein kinase whose accumulation is controlled by progesterone and occurs upstream of Cdk1^11,32^ but whose activation depends on Cdk1 activity and is involved in spindle assembly and Plk1 activation^7,8^. MELK, initially described as Eg3, is a protein kinase that accumulates in mitosis in somatic cells^36^ and whose mRNA undergoes polyadenylation during meiotic divisions, suggesting translational regulation^36^. Dnmt1 is a DNA methyltransferase that controls DNA re-methylation after S-phase^37^. Interestingly, strong accumulation of Dnmt1 was also detected by mass spectrometry in human oocytes (Supp. Fig. 8)^38^.

To validate our approach, we used as a positive control a well-characterized protein, Speedy/RINGO, which directly binds and activates Cdk1^16,17,39^. Speedy is translated in prophase-arrested oocytes but undergoes constant degradation; progesterone inhibits this degradation, leading to its accumulation^39^, that we also detect in our mass spectrometry dataset (Fig. 2B). Using our translation reporter assay, we found that Speedy translation is already relatively high in prophase-arrested oocytes and does not significantly increase upstream of Cdk1 activation (Fig. 4A-B). Speedy translation is only moderately activated downstream Cdk1 activation (Fig. 4A-B). These results support the model that progesterone triggers Speedy accumulation by promoting its stabilization through inhibition of degradation, but not by stimulating its translation, before Cdk1 activation. Applying the same translation reporter approach, we observed that the translation of MELK and Aurora-A is modest in prophase oocytes and remains unchanged upstream of Cdk1 activation after progesterone treatment, indicating that they are regulated similarly to Speedy, primarily at the level of protein stability (Fig. 4C-F). In contrast, Dnmt1 translation is strongly repressed in prophase but is reproducibly induced by progesterone both upstream and downstream Cdk1 activation (Fig. 4G-H), resembling the translational control previously described for Mos^11^. Together, these results demonstrate that both translational activation and protein stabilization contribute to the accumulation of specific proteins in response to progesterone and that proteins under either regulatory control can be detected by our proteomic approach.

Interestingly, both Dnmt1 and MELK accumulate in 1000µm oocytes in response to progesterone but not in 800 µm (Fig. 4I-K), suggesting that both translation-dependent and - independent mechanisms are acquired during the growth phase from 800µm to 1000µm.

## Discussion

Our study reveals how the molecular capacity to activate Cdk1 is progressively acquired during oogenesis. We demonstrate that oocytes first acquire the ability to accumulate proteins in response to progesterone, independently of Cdk1 activation. Indeed, following hormonal stimulation, 1000µm oocytes remodel their proteome in a manner similar to fully-grown oocytes, despite their inability to activate Cdk1. By combining Cip1-mediated inhibition of Cdk1 with TMT-based quantitative proteomics and functional assays, we delineate these upstream proteome changes, which occur downstream of PKA inhibition but independently of Cdk1 activation.

### The acquisition of meiotic competence is stepwise and developmentally regulated

Our findings reveal that the capacity to resume meiosis is acquired in a stepwise manner during oogenesis (Fig. 5). This developmental regulation likely safeguards against premature meiotic resumption in small oocytes, which would result in poor-quality eggs and embryos with limited developmental potential. We demonstrate that oocytes around 1000 µm in diameter already respond to progesterone by modulating protein homeostasis and accumulating upstream regulators such as Cyclin B1 and Mos. However, they fail to undergo NEBD. This failure is associated with insufficient levels of multiple components of the Cdk1 autoamplification loop, as Plk1, Bora and Aurora-A. Importantly, co-expression of Plk1, Bora and Aurora A rescues progesterone-induced Cdk1 Y15 dephosphorylation in 1000µm oocytes. This result demonstrates that the limited abundance of these proteins directly contributes to the failure of small oocytes to activate Cdk1. However, Cdk1 Y15 phosphorylation remains higher than in fully-grown oocytes, indicating that additional limiting components might be required to achieve full activation of the Cdk1.

These observations suggest that the acquisition of meiotic competence proceeds through at least two discrete steps: first, the ability to modulate translation and protein turnover in response to progesterone; second, the accumulation of some critical components of the autoamplification-loop that ensures robust Cdk1 activation. Our proteomic comparison of 800µm and 1000µm oocytes was limited by the number of samples that could be analyzed together in a single TMT experiment, precluding a detailed statistical analysis of this transition. Nonetheless, we speculate that 800µm oocytes may lack activators, such as RNA-binding proteins, required to stimulate translation downstream of PKA inhibition. Alternatively, they may express high levels of translational inhibitors that are insensitive to PKA downregulation. Interestingly, CPEB1, a key translational regulator degraded only after Cdk1 activation, is more abundant in 800-µm oocytes and remains stable following progesterone treatment, potentially contributing to the blockade of downstream translation in these oocytes.

A key question emerging from our findings is how the components of the autoamplification loop themselves accumulate during oocyte growth. Although fully-grown oocytes are transcriptionally quiescent, low levels of transcription may persist in 1000µm oocytes, raising the possibility that some of these components are newly transcribed during this transitional phase. Alternatively, their accumulation may rely on translational activation or stabilization of proteins produced from pre-existing mRNAs. Dissecting the transcriptional, post-transcriptional and post-translational mechanisms controlling each component will help clarifying the architecture of the meiotic-competence acquisition program.

### Protein homeostasis is regulated both at the translational and proteolytic levels

We show that the accumulation of key Cdk1 activators is controlled not only by *de novo* translation but also by reduced protein turnover. This observation extends our previous findings on Cyclin B1, which accumulates in response to progesterone not through translational upregulation, but via stabilization^11^. Here, we identify additional proteins, such as Speedy, Aurora-A, and MELK, that follow a similar regulatory logic. These proteins share the feature of having relatively high basal translation in prophase-arrested oocytes and of undergoing only modest translational induction prior to Cdk1 activation. This suggests a broader mechanism of protein stabilization operating downstream of PKA inhibition, potentially involving the regulation of E3 ubiquitin ligases. In prophase oocytes, these ligases may actively degrade a set of Cdk1 activators. Upon PKA inhibition, their inactivation could lead to rapid protein stabilization and accumulation. This model would explain the coordinated increase of several upstream regulators and shifts the focus from individual targets to a possible shared degradative pathway. Identifying the responsible ubiquitin ligases and understanding their regulation by PKA will be an important avenue for future research.

Altogether, our study defines a new proteomic and functional framework for understanding how protein accumulation upstream of Cdk1 activation regulates the ability and the timing of meiotic resumption. It identifies new key regulators of this process, highlights the coordinated regulation of protein stability and translation, and reveals how the acquisition of meiotic competence is tightly coupled to oocyte growth. These findings extend beyond the field of oogenesis. By shedding light on the molecular mechanisms linking cell size to the regulation of cell division, they open new avenues for investigating the checkpoint pathways that coordinate cell growth and cell-cycle progression, whose dysregulation is a key contributor to tumorigenesis.

## METHODS

### Animals

Adult *Xenopus laevis* females were obtained from the Centre de Ressources Biologiques Xénopes (CNRS, France) and maintained in the aquatic animal facility of the IBPS in accordance with French legislation (Animal Facility Agreement: #A75-05-25). All procedures involving animals were reviewed and approved by the French Ministry of Higher Education and Research (authorization APAFIS #45718-2025021111055055v2).

### *Xenopus* oocyte handling

Oocytes were collected from non-hormonally primed *Xenopus laevis* females. Anesthesia was induced by immersing the animals for 30 minutes in a solution containing 1 g/L MS222 (Sigma) buffered with sodium bicarbonate. Ovarian lobes were excised and enzymatically treated in M buffer (10 mM HEPES pH 7.8, 88 mM NaCl, 1 mM KCl, 0.33 mM Ca(NO₃)₂, 0.41 mM CaCl₂, 0.82 mM MgSO₄) containing 0.4 mg/mL dispase II (Roche, 04942078001) for 3 hours, followed by 0.4 mg/mL collagenase (Sigma, C9891) for 1 hour. After digestion, oocytes were thoroughly washed with 2 liters of M buffer to remove residual enzymes. Smaller stages were selected using a calibrated lengths to allow 100μm resolution. Maturation was induced with 1 µM progesterone. Oocytes displaying the first pigment rearrangement at the animal pole were considered to have undergone NEBD.

### Plasmids, mRNA preparation, antisense oligonucleotides and recombinant protein purification

Plasmids used in this study include GST-XenCip1 (NM_001094464.1), v5-reporter-3′UTRs-Aurka, v5-reporter-3′UTRs-Spdyc, v5-reporter-3′UTRs-MELK, v5-reporter-3′UTRs-Dnmt1; MBP-GST-v5. Full plasmid sequences are available in Supplementary Table 1. Constructs were generated using overlap extension PCR or Gibson assembly and verified by Sanger sequencing. mRNAs were synthesized using the mMESSAGE mMACHINE T3 Transcription Kit (Ambion, AM1348). When specified, transcripts were polyadenylated to a length of 150–200 nucleotides using the Poly(A) Tailing Kit (Ambion, AM1350). All mRNAs were purified with the MEGAclear Kit (Ambion, AM1908). Concentration was determined via NanoDrop, and transcript integrity and polyadenylation status were confirmed by electrophoresis. The amounts of mRNA injected per oocyte were as follows: 0.5 ng for v5-reporter-3′UTRs-Aurka, v5-reporter-3′UTRs-Spdyc, v5-reporter-3′UTRs-MELK, v5-reporter-3′UTRs-Dnmt1; 0.1 ng for MBP-GST-flag-HA-v5 (polyadenylated). Recombinant proteins were expressed and purified using glutathione-agarose columns (Sigma, G4510) for GST fusions. The injected amounts per oocyte were: 60 ng of GST-XenCip1-ORF. 100ng mix of two antisense sequences were injected in each oocytes. Antisense oligonucleotides were designed according to ^40^, and BLASTed against the *Xenopus leavis* genome to reduce the risk of aspecific binding. Melk antisense sequences: CTGTTTCATGCAGCTCATAG (AS1), ACTTTAGCAAATCCACCTG (AS2). Dnmt1 antisense sequences: GCCGTTTCCTGACATCAG (AS1), CACGGGCTGGTGATTT (AS2).

### Oocyte lysates, SDS-PAGE, and western blotting

Oocytes were lysed in 10 µL of extraction buffer (80 mM β-glycerophosphate pH 7.3, 20 mM EGTA, 15 mM MgCl₂) per oocyte. Proteins were resolved on 10%, 12%, or 15% SDS-PAGE gels and transferred onto nitrocellulose membranes (Amersham, 10600015) using a semi-dry blotting apparatus. Primary antibodies were diluted in PBS containing 3% BSA and 0.05% NaN₃. The following antibodies were used: anti-Karyopherin (goat, 1:2000, Santa Cruz sc-1863), anti-CPEB1 (mouse, 1:2000)^41^, anti-CyclinB1 (goat, 1:2000)^11^, anti-CyclinB2 (mouse, 1:1000, Abcam ab18250), anti-dnmt1 (1:1000, (D63A6) cell signalling), anti-melk (1/1000)^42^, anti-Mos (rabbit, 1:500, Santa Cruz sc-86), anti-pMAPK (mouse, 1:2000, Cell Signaling 9106), anti-Erk1/2 (rabbit, 1:2000, Santa Cruz C-16 and C-14), anti-pT320-PP1 (rabbit, 1:1000 for endogenous detection and 1:30000 for kinase assay, Abcam ab62334), anti-pY15-Cdk1 (rabbit, 1:1000, Cell Signaling 9111), anti-Cdk1 (mouse, 1:5000, Invitrogen MA-91598), anti-V5 (mouse, 1:1000, Invitrogen 46-0705), anti-pT210-Plk1 (1:1000, Abcam Ab39068), anti-pS216-Cdc25 (1:2000, Cell Signaling 4901S), and anti-Aurora A (rabbit, 1:1000). HRP-conjugated secondary antibodies (Jackson ImmunoResearch) were used at 1:10000.

### Mass spectrometry sample preparation

Ten to twenty oocytes were flash-frozen in liquid nitrogen and lysed as described previously^43^ in freshly prepared lysis buffer (1 mL per 20 oocytes: 250 mM sucrose, 1% NP-40, 10 mM EDTA, 25 mM HEPES pH 7.2, 10 µM cytochalasin D, 10 µM combretastatin, and Complete Roche Mini protease inhibitor tablet, 1 per 10 mL). Oocytes were lysed by pipetting up and down 20 times with a P1000, incubated at 4 °C for 10 min, vortexed for 10 s, and centrifuged at 2,500 × g for 4 min at 4 °C to pellet yolk. Samples were gently flicked 10 times to resuspend proteins associated with surface lipids, and the supernatant was collected without disturbing the yolk pellet. SDS (2%) and DTT (5 mM) were added to each sample, followed by vortexing and incubation at 60 °C for 15 min to denature proteins and reduce disulfide bonds. Cysteines were alkylated with 50 mM iodoacetamide (from a 1 M stock in anhydrous N,N-dimethylformamide) for 1 h at room temperature in the dark, and excess reagent was quenched with 30 mM DTT for 20 min. Proteins were isolated by chloroform/methanol precipitation as described previously ^44^ and resuspended at ∼2 µg/µL in 6 M guanidine hydrochloride (GuHCl), 10 mM EPPS pH 8.5. Protein concentration was determined by micro BCA assay (ThermoFisher #23235), and 30 µg of each sample was diluted to 2 M GuHCl with 10 mM EPPS pH 8.5 and digested with LysC (20 ng/µL; Wako) for 16 h at room temperature. Samples were further diluted to 0.5 M GuHCl and digested with additional LysC (20 ng/µL) and trypsin (10 ng/µL; sequencing-grade modified, Promega #V51111) for 18 h at 37 °C. Samples were dried in vacuo and resuspended in 200 mM EPPS pH 8.0 at ∼1 µg/µL. Peptides were labeled with 12 µL of TMTPro 18-plex reagent (Thermo #A44520; 20 µg/µL in anhydrous acetonitrile) for 2 h at room temperature and quenched with 6 µL of 5% hydroxylamine for 15 min (MilliporeSigma #438227). All samples were combined, dried in vacuo, acidified to pH < 1 with TFA, and desalted using a Waters Oasis HLB 30 mg cartridge (WAT094225) with three 1 mL washes of 1% formic acid followed by elution with 35% acetonitrile/1% formic acid. Samples were dried, resuspended in 150 µL of 10 mM ammonium bicarbonate pH 8.0, and clarified by ultracentrifugation at 80,000 rpm for 30 min at 4 °C in a TLA-100 rotor. 100 µL were fractionated by reverse-phase chromatography as described previously^45^ into 24 fractions, which were desalted using StageTips. Approximately 8 µg of peptides per fraction were analyzed by mass spectrometry. Peptide fractions were analyzed on a Thermo Orbitrap Eclipse Tribrid mass spectrometer equipped with a FAIMS Pro interface and an EASY-nLC 1200 HPLC. Data were acquired in data-dependent positive-ion mode using a 271 min gradient (solvent A: 0.125% formic acid, 2% DMSO; solvent B: 0.125% formic acid, 2% DMSO, 80% acetonitrile): 0–2% B (0–2 min), 2–3% B (2–12 min), 3–18% B (12–255 min), and 18–100% B (255–271 min). FAIMS Pro was operated at standard resolution with a carrier gas flow of 3.8 L/min, using four parallel experiments at compensation voltages of −32, −42, −52, and −62 V. MS1 survey scans were acquired in the Orbitrap at 120,000 resolution over 400–1100 m/z with an RF lens setting of 60%, AGC target of 1 × 10⁶, and maximum injection time of 200 ms. Precursors with charge states 2–6 and intensities ≥5 × 10⁵ were selected for MS2 using a 60 s dynamic exclusion window and 10 ppm mass tolerance. MS2 scans were acquired in the Orbitrap at 50,000 resolution following HCD fragmentation (normalized collision energy 37%), with a first mass of 110 m/z, AGC target of 5 × 10⁴, and maximum injection time of 120 ms.

### Mass spectrometry data analysis

Mass spectra were searched and quantified as described previously^46^. The Xenopus laevis v10.1 (JGI) protein reference was supplemented with mitochondrial genome sequences and used as the protein sequence database Xenopus laevis v10.1 (JGI) Genome Assembly. Searches were performed using Comet ^47^ with a precursor tolerance of 50 ppm, up to two missed cleavages, and up to three variable modifications per peptide (methionine oxidation, +15.994914 Da; asparagine deamidation, +0.984016 Da). Static modifications included carbamidomethylation of cysteine (+57.021463 Da) and TMT-Pro modification of lysine residues and peptide N-termini (+304.20660 Da). Peptide-level false discovery rate (FDR) was controlled at 1% using the target–decoy strategy^48^, and protein-level FDR was further filtered to 1% using protein picker^49^. TMT-Pro reporter ions were quantified using a 20 ppm tolerance and corrected for isotopic impurities using the manufacturer’s impurity matrix. In total, 559,536 peptides were identified, and 520,787 peptides with summed signal-to-noise >400 across all 18 conditions were quantified. Total reporter ion signal per condition was normalized to equal total intensity (all samples were within ∼20% prior to normalization). For proteins quantified by multiple peptides, only peptides with MS2 isolation specificity ≥0.98 were used, yielding 242,614 peptides mapping to 12,223 Xenopus proteins. Xenopus gene symbols were mapped to human gene symbols as described previously^50^.

### Translation reporters

Reporter experiments were performed by co-injecting 0.5 ng of v5-GST reporter and 0.1 ng of v5-MBP-GST loading control. Lysates were analyzed by western blot with anti-V5 antibody. The reporter signal was normalized to the loading control to obtain a normalized ratio, then standardized to compare different replicates using the average ratio of each replicate. Graphs and statistical analyses were carried out with Prism.

### *In vitro* Cdk1 kinase assay

Cdk1 kinase activity from oocyte extracts was assessed as described previously^51,52^.For in vitro assays using recombinant proteins, reactions were assembled in a buffer containing 20 mM HEPES, 2 mM EGTA, 10 µM β-mercaptoethanol, 1 mM ATP, 100 mM MgCl₂, 1 mM NaF, and 0.2 mg/mL Na₃VO₄. Each reaction contained 1 µg of GST-PP1 as substrate. Samples were incubated for 30 min at 30 °C.

## Acknowledgments

We thank Jean-Pierre Tassan (IGDR, CBNRS-Rennes University) for the gift of the anti-MELK antibody. This work was supported by the National Center for Scientific Research (CNRS) and Sorbonne University, the National Research Agency (ANR) grants ANR-23-CE12-0045-01 (EMD), the ARC foundation grant ARCPJA2023080006901 (EMD). LP was supported by NIH’s OD R24 grant OD031956.

## Author contributions

**Mariam Gachechiladze:** Experiments; Data Analysis; Data Curation; Writing. **Sarah B. Eivers:** Experiments; Data Analysis; Data Curation; Writing; **Robert Poulhe:** Experiments; Data Analysis. **Matthew Sonnet:** Mass spectrometry experiment and analysis, Writing. **Leon Peshkin:** Mass spectrometry experiment and analysis, Writing. **Catherine Jessus:** Data analysis, Data Curation; Writing; Supervision**. Enrico Maria Daldello:** Conceptualization; Experiments; Data Analysis; Data Curation; Writing; Supervision. **Mariam Gachechiladze** and **Sarah B. Eivers** contributed equally to this work.

